# Mind the crack: Crack-arrest holes and soft-suspension support integration in cryo-lamella preparation for improved resistance to crack formation, fracture and deformation

**DOI:** 10.64898/2026.03.05.709965

**Authors:** Sergey Gorelick, Sylvain Trépout, Sailakshmi Velamoor, Patrick Cleeve, Georg Ramm

## Abstract

Preparing electron-transparent cryo-lamellae is inherently a serial and low-throughput process. Once the lamellae are milled, these thin structures endure both mechanical and thermal stress, and as a result many valuable lamellae crack or even disintegrate entirely. This loss is often regarded as a “lamella tax”, *i.e*. an unavoidable cost of working with such fragile specimens. In this work, we introduce two modifications to the standard lamella-preparation workflow aimed at improving lamella mechanical resistance to crack formation and external stress. The first modification involves milling arrays of perforations directly within the lamella body. These perforations are designed to function as crack-arrest holes, intercepting cracks as they appear and preventing, or at least delaying their further propagation. By slowing crack growth, these features increase the likelihood that the lamella remains intact long enough to complete cryo-TEM imaging. The second modification replaces the conventional rigid attachment of the lamella to the surrounding cellular bulk material with a softer suspension using ring-shaped springs formed by ion beam milling. Mounting the lamella on smooth annular springs provides mechanical compliance both across and along the lamella axis, as well as at intermediate angles and in the out-of-plane direction. This flexibility allows the lamella to accommodate larger stresses and deformations without reaching its mechanical failure threshold. We fabricated a series of test lamellae incorporating different crack-arrest hole geometries, as well as lamellae suspended on soft annular springs. We performed high-resolution cryo-TEM imaging to characterise the perforations themselves and characterised the captured crack geometry within the lamellae at the highest level of detail achieved to date. TEM imaging shows crack interception and guided, non-catastrophic failure paths, while simulations confirm lowered stress in suspended lamellae.

---

Cryogenic focused ion beam (cryo-FIB) milling is a well-established method for creating electron-transparent samples for cryo-electron tomography (cryo-ET). In most workflows, cells are placed or grown on electron microscopy (EM) grids, which are then plunge-frozen. Thin cryo-lamellae are produced directly on these frozen grids using cryo-FIB milling [1, 2, 3, 4, 5, 6]. The resulting lamellae, usually thinner than 200-nm, are transferred into a cryo-transmission electron microscope (cryo-TEM) for high-resolution imaging of cells in a near-native state. After milling, the lamellae are exposed to several types of stress, including mechanical handling and changes in temperature. During the cryo-transfer between the FIB instrument and the cryo-TEM, the lamellae can undergo temperature shifts of several tens of degrees, which increases their vulnerability to damage. These temperature shifts, together with the mechanical forces involved in handling the grids, can make the lamellae structurally unstable. Because the lamellae are so delicate, even small deformations can be detrimental. In many cases, the combination of mechanical and thermal stresses is enough to cause a lamella to crack or break. Cryo-FIB milling is also a relatively low-throughput technique, since lamellae have to be prepared one at a time. Even with automated routines, an experienced operator typically manages to prepare 15–25 lamellae in a single session. Each lamella takes roughly 30 minutes of instrument time (not counting the substantial effort required beforehand to prepare the sample). Thus, when a lamella cracks or disintegrates, the impact is substantial resulting in the loss of time, effort, and instrumentation resources, as well as a missed opportunity to acquire valuable cryo-TEM data. The mechanical stability of cryo-FIB-prepared lamellae is becoming a common topic in literature. Several studies have proposed strategies to enhance the robustness and survivability of cryo-FIB-milled lamellae. Wang et al. [7] describe a method for producing mechanically stable large-area tissue lamellae and discuss common modes of failure (see Supplementary Fig. 5d in [7]). Their approach involves disconnecting one edge of long, wide lamellae from the bulk material to prevent fracture. They also introduce a “furrow–ridge” design consisting of alternating thinned regions (furrows) and thicker supporting ribbons (ridges), which increases stiffness and reduces bending. This approach has proved effective in improving lamella stability. Kelly et al. describe the “notch milling” strategy, developed within the Waffle Method sample-preparation workflow [8, 9]. The notch separates one tip of the lamella by 200 nm from either side from the bulk of the sample without fully releasing it. This partial separation allows limited motion of the lamella, enabling some stress relief during handling and transfer. Notch milling has been reported to significantly enhance lamella survival during grid transfers and cryo-TEM imaging. In practice, however, its main advantage lies in facilitating final lamella thinning. Without the notch, internal stresses can induce lamella bending during ion-beam milling, complicating the production of uniformly thin lamellae. Improved survivability during handling is therefore a secondary but valuable benefit. Wolff et al. introduced the “micro-expansion” strategy, which involves milling small trenches on either side of a cell lamella to improve mechanical integrity [10]. Although originally intended to reduce lamella bending caused by stress propagating from the grid (also noted by Zhang et al. [11]), this method was also found to increase overall lamella survivability. Micro-expansion gaps have since become a standard and widely adopted element of lamella-preparation protocols due to their simplicity and effectiveness.

Reports of failed or fractured lamellae remain relatively rare, as publications typically highlight only successfully prepared examples. Nevertheless, available literature [7], community communications [12], and our own observations indicate that lamellae most often fracture near the points where they remain connected to the bulk material. This suggests that mechanical stresses concentrate at these edges. In conventional workflows, lamellae are milled using rectangular patterns. The focused ion beam removes material from regions above and below the area of interest, leaving a thin lamella with sharp corners at its attachment edges. These sharp internal angles inherently promote stress concentration. We have previously demonstrated that using filleted milling patterns, *i*.*e*. replacing sharp corners with smooth, rounded transitions, reduces such stress concentrations and in turn lowers the risk of lamella breakage, thereby increasing the yield of lamellae suitable for TEM [13]. Recently, Wachsmuth-Melm et al. [14] proposed a trapezoidal milling geometry. Originally designed to support dual-axis cryo-ET by reducing beam shadowing, this geometry also provides improved mechanical stability based on simulation results. The obtuse angles in the trapezoid create a smoother transition between the lamella and the surrounding bulk material, mitigating stress concentration in a manner similar to that achieved with corner fillets.

Crack formation in a lamella whether during thinning, handling, or cryo-TEM examination, is an undesirable event that can lead either to catastrophic failure and complete loss of the lamella or to significant interference with cryo-TEM imaging. Cracks in a lamella generally form at sites of elevated stress, which often arise from pre-existing defects, material inhomogeneity, or residual stresses within the lamella. Once a crack initiates, it can advance as the local stress at the crack tip concentrates external or internal forces, breaking molecular bonds and driving further fracture propagation. Surface imperfections such as FIB-induced curtaining, and material impurities can serve as stress risers, *i*.*e*. localised points of elevated stress that act as common initiation sites. Residual in-plane tensile stresses introduced during lamella fabrication can also trigger cracking even in the absence of external loading. The subsequent crack path (including tearing and branching behaviour) depends on multiple factors, such as the exact initiation site in the lamella, the propagation velocity, the associated elastic energy release, and other local mechanical conditions.

In this study, we investigate the incorporation of perforations directly into the lamella (Fig. 1a-d) to enhance its mechanical stability and improve its ability to withstand crack formation that might otherwise lead to catastrophic failure. These perforations serve two complementary functions. Firstly, they act as “micro–micro-expansion” (*µµ*-expansion) gaps [10], increasing the flexibility of the lamella and allowing it to accommodate larger in-plane (*xy*-plane in Fig. 1) and out-of-plane stresses (along the *z*-axis in Fig. 1). This added compliance helps the lamella bend and flex more without reaching its mechanical limits and fracturing. This stands in contrast to conventional lamellae that remain rigidly anchored at their edges to the bulk material and therefore tend to bend or twist in a more constrained manner. Secondly, the perforations are intended to function as crack-arrest holes, capturing cracks that initiate within the lamella and preventing or mitigating their further propagation into the lamella body. Crack-arrest holes [15, 16, 17] are a long-standing and widely used technique for slowing or stopping fatigue-crack growth in structures where the damaged structure replacement is not feasible (such as bridges or aerospace components, or even cryo-lamellae from now on). The basic method involves drilling a hole at the crack tip to blunt the sharp crack front, replacing the near-zero radius of curvature (which produces extremely high stress concentration) with a larger, rounded boundary that lowers the local stress intensity and temporarily halts further crack propagation. However, in the case of a standard drilled hole, residual tensile stresses around the hole edge frequently promote crack re-initiation on the other side of the hole [18], rendering the repair only a temporary solution. Larger-diameter holes at the end of a crack are generally more effective since they increase the crack tip radius more, greatly decreasing the applied stress intensity and delay crack re-initiation [19, 20]. Overall, crack-arrest holes offer a simple, practical means of delaying crack growth and are commonly used when full component replacement is impractical. Crack-arrest holes are traditionally introduced only after a crack has been detected, with a hole drilled at the crack tip to slow or halt its further growth. This reactive approach, however, is not feasible for cryo-lamellae. Instead, in this work we introduce crack-arrest holes pre-emptively along the lamella edges. Because cracks in lamellae typically originate near these edge regions (where stress concentrations are also highest during bending), the presence of pre-milled crack-arrest holes is intended to intercept any newly forming cracks and prevent them from propagating or branching into the lamella interior. In doing so, these features aim to extend lamella survival long enough to complete cryo-TEM data collection.

**Figure 1:**
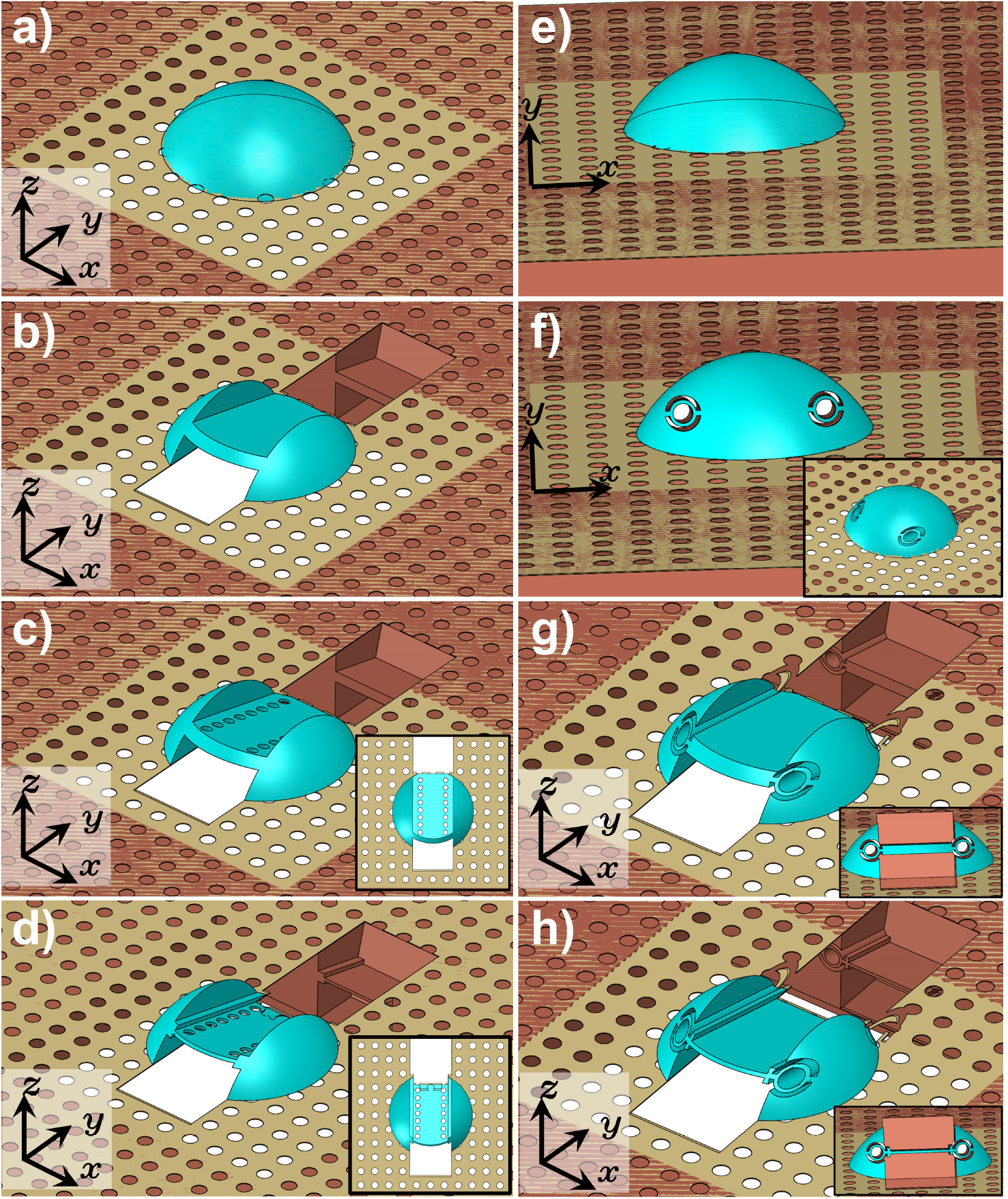
Schematic illustrating the preparation of a thin lamella from a cell on an EM grid with (a-e) crack-arrest holes and (e-h) with annular spring suspension. (a) The initial cryo-fixed cell. (b) A thick lamella is milled from the cell. (c) The stage is tilted from the lamella-milling angle used in (b) to a higher ion-incidence angle, and perforations are milled into the thick lamella. The inset shows the ion beam view for milling the perforations. (d) The stage is returned to the lamella-milling angle, and the lamella is thinned to its final thickness. (e) The initial cryo-fixed cell. (f) Annular springs are milled into the cell; the inset shows an alternative viewing angle. (g) A thick lamella is subsequently milled; the inset provides a view from the ion-beam direction at the lamella-milling angle. (h) The lamella is thinned to its final thickness; the inset again shows the corresponding ion-beam view at the milling angle.

As noted above, lamellae are conventionally anchored rigidly at their edges to the surrounding bulk material. In this configuration, any in-plane tensile or compressive stresses that develop within the lamella can cause it to bend or even buckle. To address this limitation, we propose mounting the lamella on a softer, spring-like suspension that can absorb internal stresses and also buffer the lamella against external mechanical shocks (Fig. 1e-h). Specifically, we introduce ring-shaped springs milled into the bulk material on either side of the lamella, provided that the cell volume is sufficient to accommodate them. Such annular springs offer the advantage of an angle-dependent spring constant [21, 22], meaning they provide compliant support for deformations along the *x*-axis of the lamella, the *y*-axis of the lamella (see Fig. 1 for corresponding axis), and any intermediate direction. In addition, the ring-spring geometry can accommodate out-of-plane deflections (*z*-axis), thereby enhancing the overall mechanical resilience of the lamella. The annular spring geometry provides an important advantage. Its continuous, rounded shape naturally minimises sharp corners, which occur only at the points where the spring attaches to the substrate and to the lamella. Rounded features are mechanically beneficial because they reduce stress concentration and are less likely to to contribute to elevated stress. In contrast, meander-type springs consist of multiple straight beam segments connected at sharp angles, each of which can introduce local stress concentrations and increase the risk of structural failure. The continuous, smooth curvature of the annular spring makes it a mechanically more robust option for supporting fragile lamellae.

Cracks in lamellae are most commonly observed along their edges and, in some cases, extend or branch into the central regions (Fig. A.5). However, this pattern may reflect a form of survivor bias: only lamellae with benign cracks and that remain partially intact can be inspected, while those that fail catastrophically leave no clear evidence of where the initial crack originated. For lamellae that completely disintegrate, it is therefore impossible to determine whether the fracture began at the edges or from within the lamella interior, making it difficult to place crack-arrest holes precisely at the most critical locations. It is plausible that some shattered lamellae fail due to cracks initiating in the central region, potentially triggered by excessive bending. When a lamella bends beyond its mechanical limit, it may fragment into multiple pieces. This failure mode is similar to that of how a bent spaghetti strand breaks into three or more fragments [23]. In this phenomenon, the initial break releases stored elastic energy that generates snap-back vibrations, producing secondary fractures. A similar cascading failure could occur in over-stressed lamellae.

In this study, we positioned crack-arrest holes along the sides of the test lamellae. As illustrated schematically in Fig. 2a, in the absence of such perforations a crack that initiates at either the top or bottom edge can propagate across the lamella and reach the opposite side. During this process, the primary crack may branch into secondary cracks that extend toward the lamella interior (Fig. A.5). This outcome is undesirable because it can interfere with cryo-TEM imaging, render the lamella completely unusable or even cause the lamella to break completely. By placing an array of perforations pre-emptively along the lamella edges, we aim to intercept cracks that originate and propagate along these regions (Fig. 2b). A crack-arrest hole can temporarily halt crack growth, extending the lifetime of the lamella to complete TEM data collection. Although the crack will eventually re-initiate on the other side of the hole, the delay may be sufficient to finish imaging, or the next perforation in the array may again arrest the crack and further prolong the lamella’s usable lifetime (Fig. 2c).

**Figure 2:**
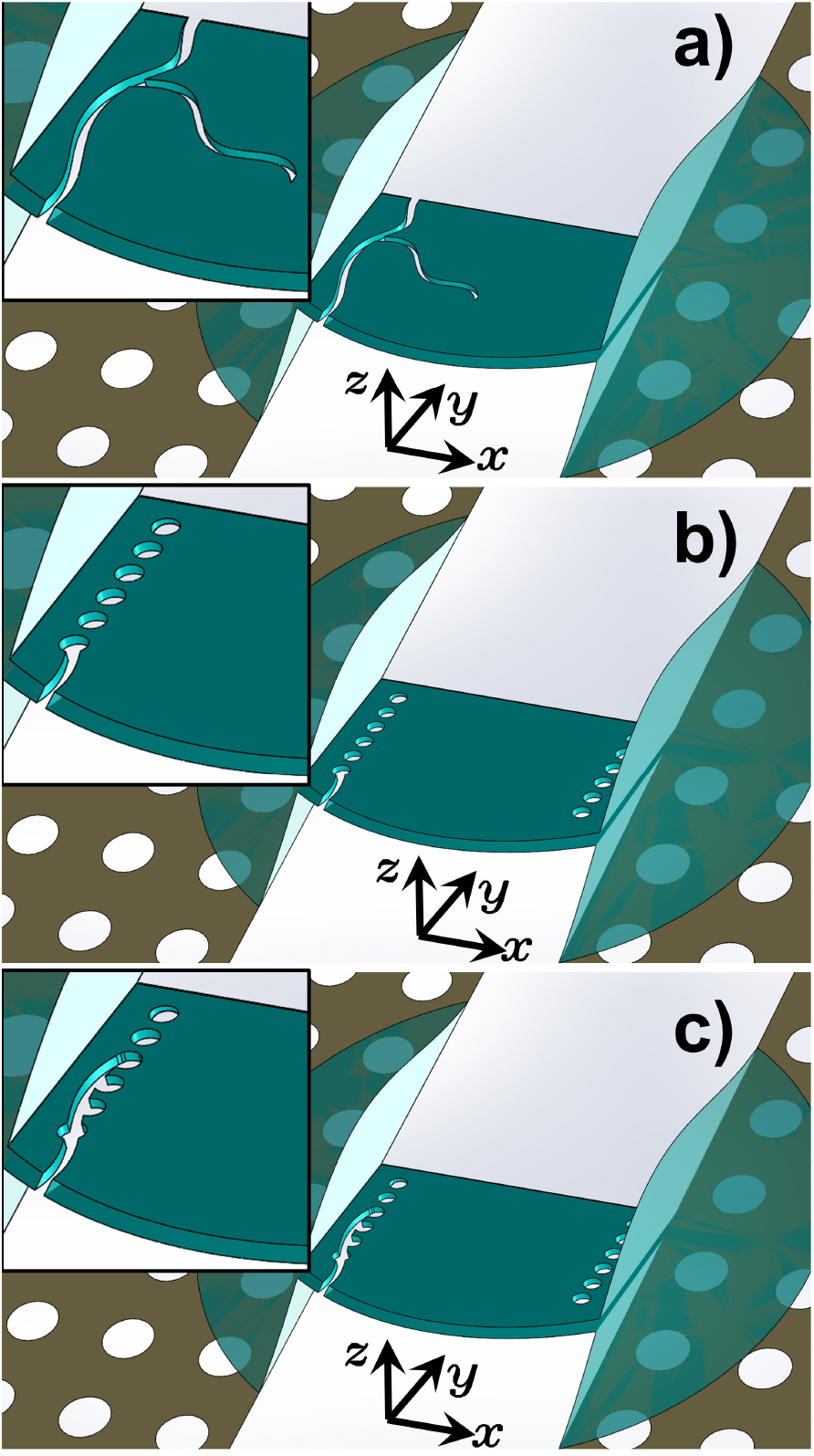
Schematic illustrating the function of crack-arrest holes. (a) A crack forms at the lamella edge, propagates along the boundary, and branches into the lamella interior. (b) In the same lamella, arrays of perforations are milled along the edges. A crack originating at the boundary begins to propagate but is intercepted by a crack-arrest hole. (c) Although a crack-arrest hole can only delay subsequent re-initiation of crack growth, the next hole in the array captures the crack again, further slowing its progression.

While covering the entire lamella with perforations would improve crack-arrest performance, it would also remove too much cellular material and reduce the area available for TEM imaging. For this reason, a balance must be found between the number of holes and their size so that the lamella remains both mechanically robust and imageable. Although placing holes along the central axis of the lamella might also be useful, in this study we focus only on edge-mounted perforations to keep the experiment simple. We test several hole sizes, recognising that larger holes are better at stopping cracks [19, 20], whereas smaller ones preserve more usable imaging area.

Figure 3 shows two lamellae prepared with arrays of side perforations: the lamella on the left contains larger holes, approximately 1.2 *µ*m in diameter, whereas the lamella on the right features smaller 0.5 *µ*m perforations. The crack-arrest holes were spaced along the *X*-direction to match the final width of the lamella, while the pitch in the *Y*-direction was set to 2 *µ*m. Because the holes were milled with the lamella tilted away from the standard milling angle—resulting in a higher ion-beam incidence on the lamella surface—the perforations are sloped relative to the lamella’s *Z*-axis. Consequently, the apparent *Y*-direction spacing appears larger due to the projection of the hole array onto the lamella surface. Fig. A.6(a,b) in Appendix A shows high-resolution electron tomograms of crack-arrest holes acquired at several stage tilts and for several perforation geometries. In our study, we were able to document several cracks within lamellae that did not lead to complete structural failure (Fig. A.6c,d). Interestingly, these cracks were typically observed in lamellae of reduced quality such as those exhibiting pronounced curtaining, surface roughness, or thickness non-uniformity (Fig. A.6e,e’).

**Figure 3:**
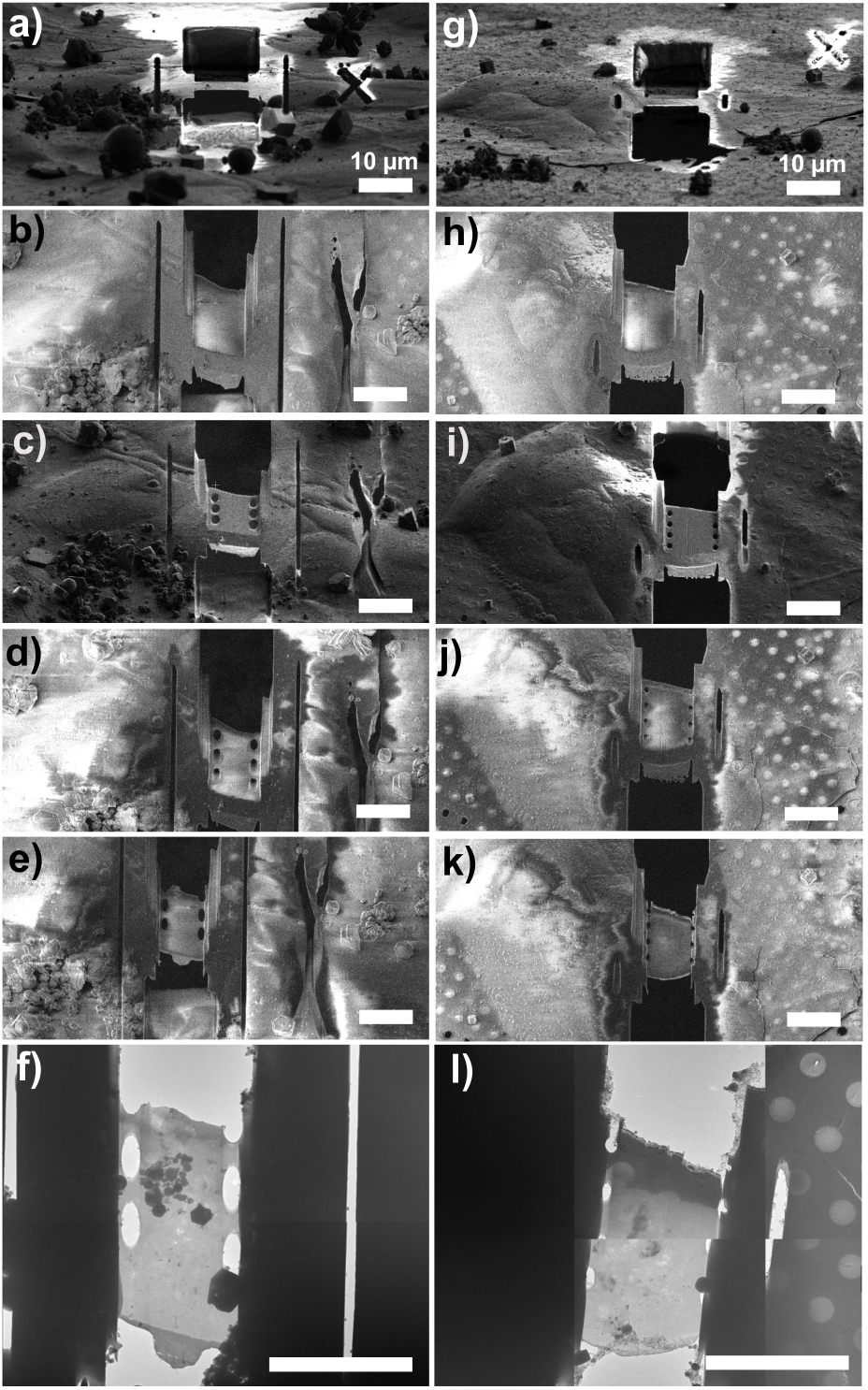
Lamellae with periodic near-edge perforations designed to function as crack-arrest holes. (a–f) Lamellae with larger perforations (1.2 *µ*m diameter) spaced along the *Y*-axis; (g–l) lamellae with smaller perforations (0.5 *µ*m diameter). (a,g) Focused ion beam views of the initial ∼ 1.5 *µ*m-thick lamellae prepared in the cells. (b,h) Corresponding SEM views. (c,i) The stage is tilted from the lamella-milling angle of 14^◦^ (lamella at 12^◦^ relative to the grid plane) to 40^◦^ and 45^◦^, respectively, and the perforations are milled into the lamellae. (d,j) After perforation milling, the stage is returned to the lamella-milling angle and SEM images are acquired. (e,k) The lamellae are subsequently thinned to their final thickness. (f,l) Corresponding low-magnification TEM images.

In one example, a crack initiated at the lamella edge and propagated along the array of crack-arrest holes (Fig. A.6e,e’). Whether the perforations contributed to initiating the crack or instead diverted its path and ultimately preserved the lamella cannot be determined, as a true control experiment is not feasible. In this case, the crack re-initiated at the opposite side of the hole, continued toward the next perforation, and exited on the other side before progressing to the far edge of the lamella consistent with the expected overall performance of simple crack-arrest holes [18].

In another case, a lamella with crack-arrest holes on both sides appeared to be of good quality immediately after milling (Fig. A.6f,f’). However, once transferred into the TEM, we observed that one side of the cellular material and hence one of the lamella’s anchoring edges was missing. The fracture followed the line of perforations, yet the majority of the lamella remained intact despite the loss of one anchor edge. Although we cannot provide definitive evidence, we speculate based on (i) the lamella’s intact appearance after milling, and (ii) the fact that a portion of the cell snapped off precisely along the perforation line, that significant internal stress developed within the lamella, likely due to cracking or deformation of the supporting grid. Under normal circumstances, such stress would be expected to cause the lamella to shatter into multiple fragments. Instead, the additional compliance introduced by the perforations appears to have guided the breakage along a controlled path, preserving the remainder of the lamella. As a result, rather than losing the specimen entirely, we were able to image nearly the full remaining surface.

Although the exact initiation point and propagation path of a crack cannot be predicted, it is still advantageous to position arrays of crack-arrest holes on the lamella in advance. Placing these perforations along the lamella edges is practical, as it minimises interference with imaging. Previous studies have shown that crack-arrest holes can influence crack trajectories and may even suppress propagation between adjacent holes due to the development of compressive stresses in the region between them [24]. These compressive stresses can reduce the driving force for crack growth, thereby helping to limit or delay further propagation.

However, while crack-arrest holes help manage the consequences of stress by interrupting crack propagation, they do not address the underlying sources of stress within the lamella. To mitigate these stresses directly, we next explore supporting the lamella on compliant, ring-shaped springs rather than leaving it rigidly anchored to the surrounding cellular bulk (Fig. 4). In this approach, the lamella is suspended on hollow, annular springs milled into the adjacent cell or tissue from which the lamella is prepared. By removing the rigid structural connection on both sides and replacing it with a smooth, compliant support, this design provides a pathway for internal stresses to dissipate instead of accumulating within the lamella. Finite-element simulations comparing a conventional lamella with one supported by a ring-shaped spring indicate that the suspended configuration experiences a noticeable reduction in internal stress (Fig. A.7a,b), albeit accompanied by a modest increase in overall deformation. The choice of annular springs is deliberate. Although many spring geometries exist, the ring-shaped design offers compliance in multiple deformation directions (in addition to the ease of implementation in milling). It can accommodate in-plane stresses and displacements allowing the lamella to shift along the *X*-axis (Fig. A.7c), while being especially compliant to shear-like, rotational deformation along the *Z*-axis (Fig. A.7d). This enhanced compliance enables the suspension to absorb out-of-plane stresses and deformations associated with lamella bending or buckling, thereby improving the overall mechanical resilience of the specimen.

**Figure 4:**
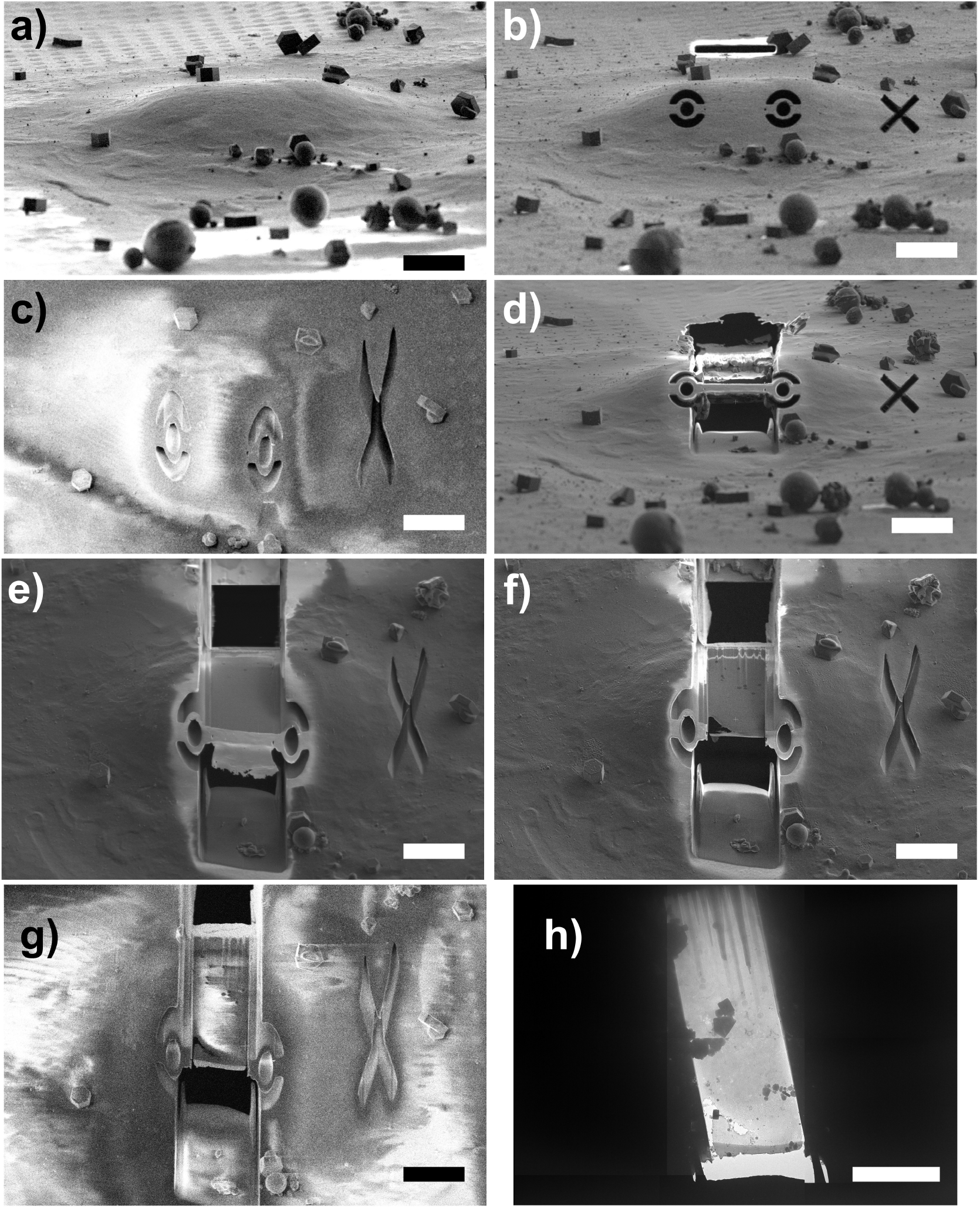
Electron micrographs of a lamella supported by annular, ring-shaped suspension springs. (a) Initial cryo-fixed cell, shown in the focused-ion-beam (FIB) view at the lamella-milling angle. (b) Ring-shaped springs and anchoring regions connecting the future lamella to the surrounding cell material are created by ion-beam milling. (c) Corresponding SEM view. (d) A thick lamella is subsequently milled, shown in the FIB view at the milling angle. (e) FIB view acquired with the stage tilted to 45^◦^. (f) FIB view of the lamella after thinning to its final thickness, also at a 45^◦^ stage tilt. (g) SEM image of the thinned lamella with the stage returned to a 14^◦^ tilt (lamella-milling angle). (h) Corresponding low-resolution TEM image of the final lamella suspended on the annular springs.

**Figure A.5:**
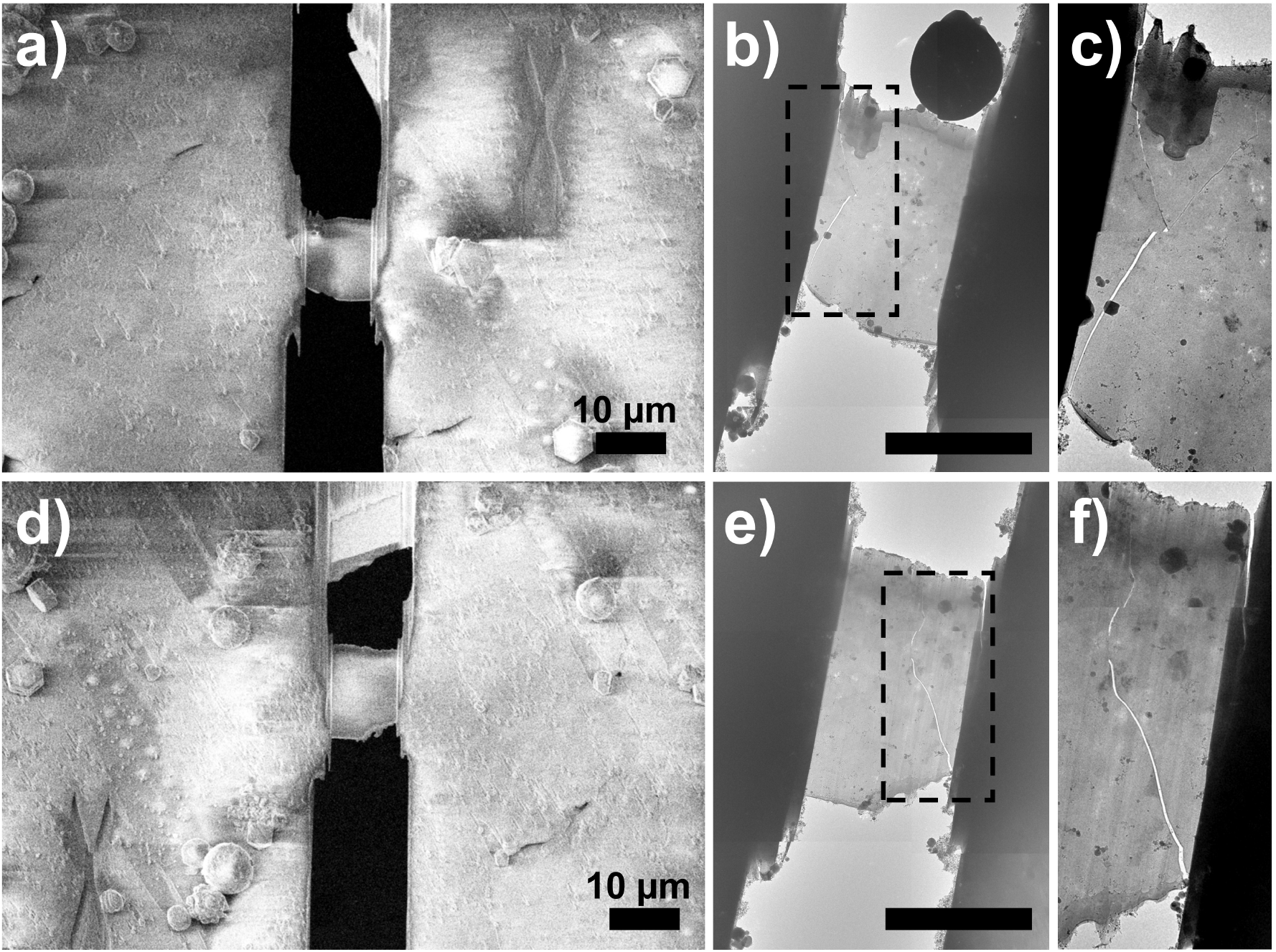
Cracks observed in lamellae after transfer into the TEM. Some lamellae were prepared without micro-expansion cuts, increasing the likelihood of fracture and providing an opportunity to examine crack formation. (a,d) SEM images of lamellae after thinning to their final thickness. (b,e) Corresponding low-resolution TEM images of the lamellae shown in (a) and (d); in both cases, cracks developed along the lamella edges. (c,f) Enlarged views highlighting the regions containing the cracks.

**Figure A.6:**
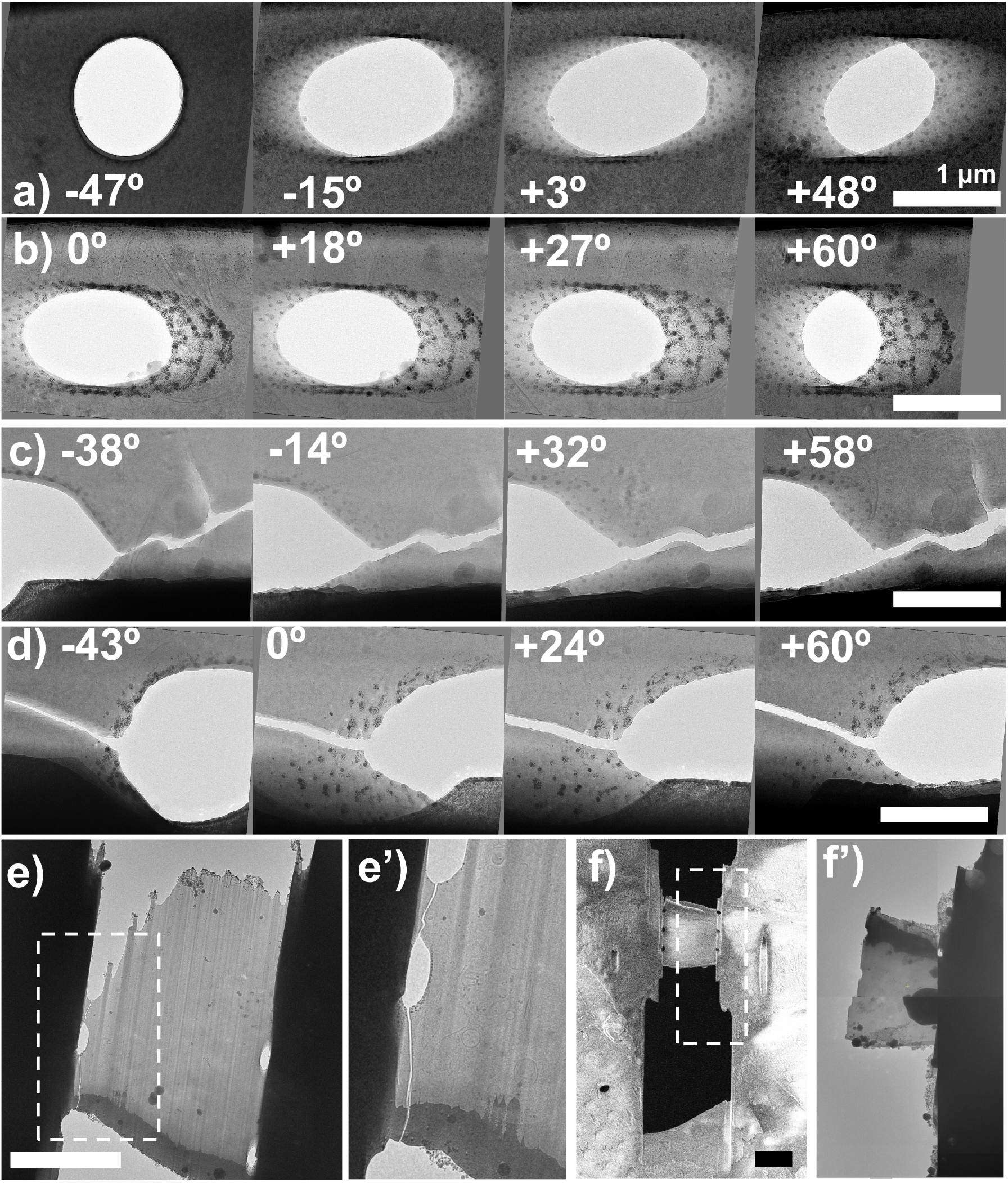
TEM imaging of crack-arrest holes in lamellae. (a,b) High-resolution tomograms of two different perforations acquired at varying stage tilts. In some cases, traces of Pt are visible inside the holes; this occurs because the perforations were milled while the lamellae were still relatively thick, allowing redeposition of sputtered protective Pt during subsequent ion-beam milling. (c,d) High-resolution tomograms of two additional perforations, imaged at different tilts, showing cracks that developed within the lamella and were intercepted at the edges of the holes. (e,e’) TEM image of a lamella with a crack along its side, accompanied by a magnified view of the affected region. (f,f’) SEM image of a lamella containing perforation, together with the corresponding TEM image demonstrating that the lamella fractured cleanly along the perforation line, leaving the remainder of the lamella intact.

**Figure A.7:**
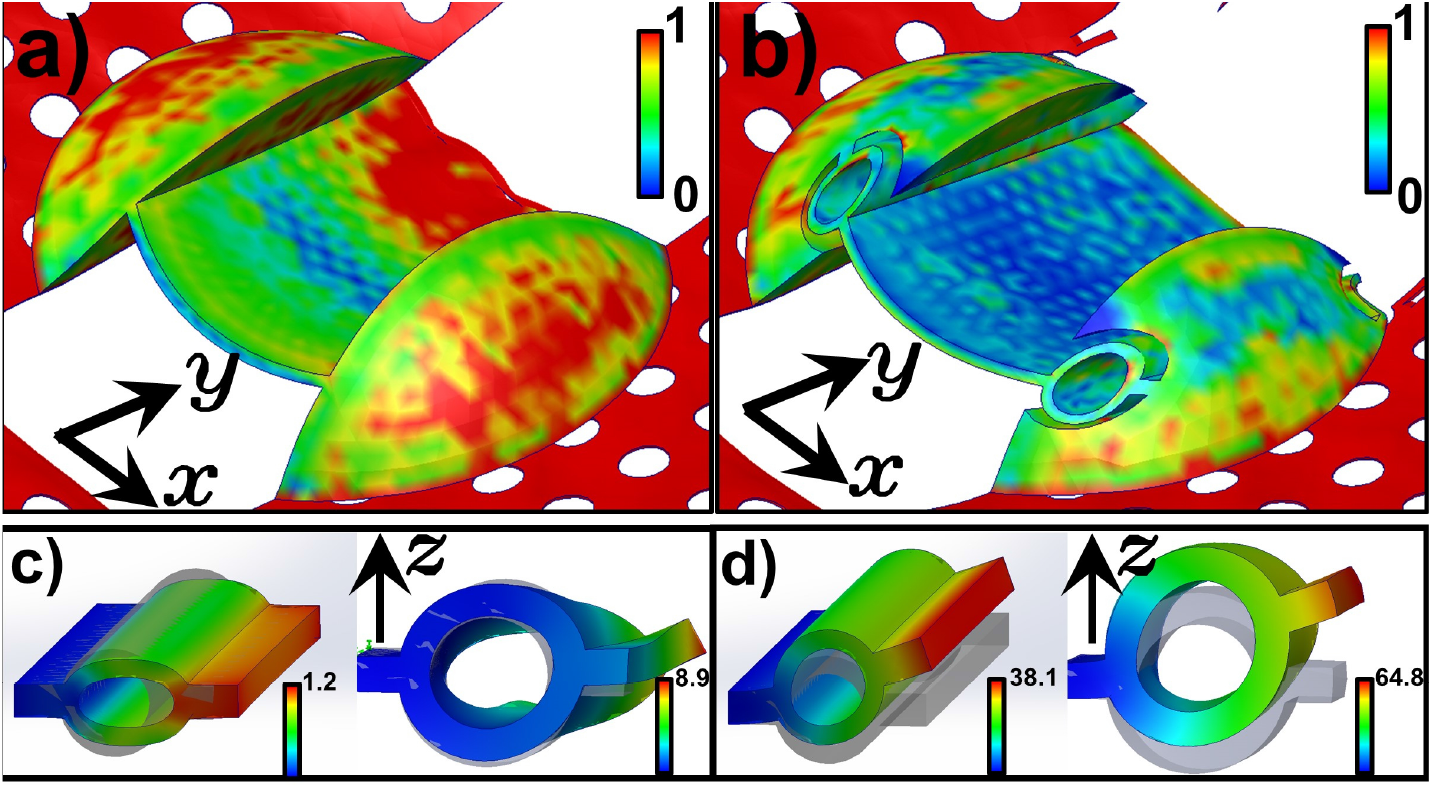
FEM simulations of a lamella supported by a ring-shaped suspension. (a) Von Mises stress distribution in a lamella subjected to a 20 K temperature drop, simulating transfer from a 100 K microscope stage into liquid nitrogen (80 K). Stress values are shown in arbitrary units for comparison. (b) Corresponding simulation for a lamella suspended on annular springs, demonstrating a substantial reduction in von Mises stress across the lamella. (c) FEM analysis of the annular spring’s deformation under an *X*-direction load, used to determine the in-plane spring constant. The simplified spring geometry is shown on the left, and a more realistic sloped and tilted design—matching the structure in panel (b) is shown on the right. (d) FEM analysis of the out-of-plane (*Z*-direction) spring constant for simplified (left) and realistic (right) spring geometries. In both cases, the applied force is 1 *µ*N, and the resulting deformations are colour-coded in nm.

To quantify the performance of the ring-shaped suspension, we note that although analytical expressions exist for estimating spring constants and elastic responses of idealised geometries, such as cylindrical shells [25], thin plates (approximating lamellae), or doubly clamped thin cantilever beams under various loads, these formulas become unreliable when the geometry includes complexities such as sloped or tapered cylindrical spring structures. For this reason, FEM (Finite Element Modeling) provides a more accurate assessment of the mechanical behaviour of our suspension design. More details on the simulations can be found in Refs. [13, 14]. FEM simulations of both simplified and more realistic spring geometries yield an in-plane (*X*-direction) spring constant of approximately 800 N/m. While this value is relatively high, it still allows the lamella some lateral motion, which would have been severely restricted under the much stiffer, fully rigid connection to the cellular bulk. More critically, the out-of-plane stiffness of the ring suspension—governing the lamella’s ability to move along the *Z*-axis (and thereby reduce bending load) is substantially lower, at roughly 30 N/m. This provides an important compliance pathway for vertical motion, enabling the lamella to accommodate deformation without accumulating damaging stress. For comparison, the bending stiffness of the lamella itself under a uniform surface load (analogous to a pressure) is even lower, around 5 N/m, making it highly susceptible to bending. Suspending this thin sheet of vitrified material on compliant springs therefore helps redistribute stresses away from the most vulnerable regions of the lamella, increasing the likelihood of its survival when subjected to deformation or external mechanical disturbances.

In summary, we introduce two modifications to the lamella-preparation workflow, namely arrays of crack-arrest holes and compliant ring-shaped suspensions. Used together with other strategies (microexpansions or notch, corner fillets), these strategies address both major aspects of lamella failure: they constrain crack propagation once it begins and reduce the internal stress build-up that predisposes lamellae to fracture. Both approaches are straightforward to implement, compatible with automated milling routines, and increase the likelihood that lamellae remain intact long enough to acquire high-quality cryo-TEM data. Given the inherently low throughput of cryo-FIB preparation, even modest improvements in lamella survivability translate into meaningful gains in experimental yield. We anticipate that the underlying design principles, such as stress-aware geometry, pre-emptive crack management, and compliant mechanical support, will be broadly applicable across cryo-ET sample preparation and will inspire further mechanically informed advances in lamella engineering.

## Materials and methods

### Cell culture on EM grids and freezing

Two types of cells were cultured for this study. Firstly, mouse embryonic fibroblasts (MEF) were maintained at 37 ^◦^C, 5% CO_2_ in in-house made DMEM High Glucose (DMEM) supplemented with 10% FBS and GlutaMAX™ Supplement (Gibco #35050061). The EM grids (Quantifoil Au R2/2) were coated with ∼ 3 nm of carbon using a Leica AF200 (Leica Systems), and glow-discharged for 30 s at 30 mA and 0.23–0.26 mBar pressure using a Pelco EasyGlow™. The cells were then seeded onto these EM grids. Samples were held in the chamber at 37 °C and 95% relative humidity prior to freezing. After 48 hours, the cells grown on the grids were transferred from the culture room to the EM facility. Freezing conditions were set up in a Leica EM-GP2 (liquid ethane held at -175 ^◦^C, chamber at 37 °C and 95% relative humidity). Samples were backside-blotted for 12 s and plunge-frozen into liquid ethane. After plunging, the grids were placed into a grid box kept in liquid nitrogen to prevent devitrification and limit ice contamination.

For testing the suspension design, we followed the same preparation protocol but used human mammalian cells as the sample material.

### Cryo-FIB milling

Lamellae fabrication was performed on a dual-beam ThermoFisher Scientific (TFS) Helios 5 UX Cryo-FIBSEM using 30 keV Ga+ beam. The grids clipped in autogrids were cryogenically transferred into the cryo-FIB, and suitable cells for thinning and milling were selected from mapping and imaging the grids with electron and ion beams. The microscope is equipped with a cryogenic shuttle with a pre-tilt of 40^◦^. The ion beam incidence with respect to the grid plane was typically set to 12^◦^ unless specified otherwise (stage tilt between 14^◦^), and kept constant throughout the lamella milling. We ensured that each selected milling site was at the eucentric point of the microscope, *e*.*g*., the point of coincidence between the electron and ion beams. In this point the sample is at 4 mm from the electron microscope’s polepiece. Next, the sample was lowered to 5.5 mm from the electron polepiece and a layer of organometallic platinum was deposited onto the grids using gas injection system (GIS) operated at 29 ^◦^C for 12 s. The lamellae milling was aided with AutoLamella (fibsemOS) software [26, 27]. Firstly, we translated the stage from one lamella site to the next, milling fiducial markers at each site to facilitate future alignments. The fiducial marks were milled using Rectangle milling pattern. Next, we proceeded to each lamella site, aligning using the previously milled fiducial markers.

For lamellae incorporating the ring-shaped suspensions, we milled the corresponding annular patterns during this stage using a 0.44 nA beam current (see Figure 1-right). The milling pattern design was implemented via the fibsemOS plugin system under the name *Circular Suspension*. This pattern is constructed using standard milling patterns available on our microscope, such as circular patterns with a non-zero inner radius and exclusion zones that prevent milling in specified regions, to define rectangular anchors to both the bulk material and the lamella. This demonstrates the flexibility of fibsemOS, which enables rapid prototyping and integration of non-standard composite milling patterns into experimental workflows.

Before the rough milling, we created micro-expansion gaps using a 2.6 nA beam current (not for the lamellae mounted on ring-suspensions). The microexpansion gaps were milled using Rectangle milling pattern. The microexpansion gaps were positioned on either side of the lamella 10 *µ*m from its centre.

We then performed rough and intermediate milling with beam currents of 2.6 nA and 0.44 nA, respectively, to define the initial, relatively thick lamellae. The rough milling step aimed at creating a 5-*µ*m-thick and 14-*µ*m-wide lamella, while the intermediate milling step aimed at creating a 1.5-*µ*m-thick and 12-*µ*m-wide lamella.

After completing the rough and intermediate milling steps, we introduced arrays of perforations into selected lamellae, intended to function as crack-arrest holes. This step was performed prior to final polishing because milling large perforations into an already thinned lamella risks bending or damaging it and can also lead to surface contamination due to redeposition of sputtered material. In principle, one could rotate and tilt the stage to orient the ion beam normal to the lamella surface for this milling step. However, to minimise stage drift, we instead tilted the stage from the lamella-milling angle to a position where the electron beam was normal to the grid surface (a 40^◦^ stage tilt, corresponding to a 38^◦^ ion-beam angle relative to the grid plane). Tilting alone (rather than rotating the stage by 180^◦^ and then tilting) also makes it easier to keep the lamella in view and accurately track it to position the perforation milling patterns. When the stage is at the standard lamella-milling angle of 14^◦^ (where the ion beam is parallel to the lamella plane), tilting the stage to 40^◦^ results in a 26^◦^ ion-beam incidence relative to the lamella plane (64^◦^ with respect to the surface normal). In some cases, we tilted the stage further, to 45^◦^ (31^◦^ incidence on the lamella plane) or even to 52^◦^ (38^◦^ incidence). During oblique milling, the perforations produced within the thick lamella naturally acquire a sloped cylinder profile. However, as the lamella is subsequently thinned from approximately 1.5 *µ*m down to below 200 nm, the height of these cylinders decreases, and the sloped geometry becomes less pronounced. A slight inclination of the hole walls remains, but it is far less noticeable in the final, thinned lamella. The milling of 200–500 nm perforations was performed using 90 pA beam current. Perforations were spaced 2 *µ*m in the *Y*-direction and 8 *µ*m in the *X*-direction, spanning the lamella’s full length and width. For this study, we manually performed the required stage movements and milling for the crack-arrest holes. The same procedure is now available in AutoLamella (fibsemOS) via the plugin system as the task “Mill Perforation” and can be optionally carried out between automated rough and intermediate milling for selected lamellae.

The polishing of all thick lamellae was performed sequentially using a 41 pA beam current to produce lamellae with a nominal thickness of 180 nm. Rough, intermediate and final polishing steps were all performed using “cleaning crosssection” patterns.

Lamella setup was typically carried out under supervision using the Autolamella interface to ensure precise placement of milling patterns, fiducial markers, and microexpansion gaps. During this setup phase, fiducials were milled and reference images were taken for future alignments in the milling process. Once all lamella positions were defined, rough and intermediate milling were performed sequentially across all the sites. At each location, alignment was refined using the fiducial markers before proceeding with thinning to approximately 1.5 *µ*m. These stages were executed autonomously, without direct supervision. Once the rough and intermediate milling were complete, the final polishing step was conducted by revisiting each stored lamella position realigned at each position using the same fiducial markers. This stage was typically supervised to allow for fine adjustments, if needed, to the milling patterns, particularly in response to minor sample drift or grid deformation. The Autolamella has recorded the process and intermediate steps in images. All SEM images were collected at 2 keV beam energy, 50 pA beam current, 1 *µ*s dwell time and 1536 × 1024 pixels image resolution. All FIB images were acquired at 30 keV beam energy, 41 pA beam current, 1 *µ*s dwell time and 1536×1024 pixels image resolution.

### Cryo-TEM tomography

After FIB milling, EM grids were promptly (within 12 hours) transferred into a Titan Krios G4 (ThermoFischer Scientific) cryo-TEM, operating at 300 kV and equipped with a Selectris-X energy filter and a Falcon 4i direct electron camera. Very-low magnification (740×) and low magnification (3,600×) images of the cryo-FIB lamellae present in this work were collected as part of the tomography workflow using the TFS Tomography TOMO5 software. Electron-Event Representation (EER) tilt-series were collected at 53,000× magnification (corresponding pixel size is 2.39 Å) using a 20 eV slit. Images were collected between -44^◦^ and +70^◦^ stage tilts every 2-3^◦^ using a dose-symmetric scheme [28], corresponding to a ± 56^◦^ rotation around the angle at which the lamellae were milled (about +12^◦^). The total electron fluence used per tilt-series was 100-120 e^−^/Å^2^.

### Cryo-TEM alignment

EER images were motion-corrected and averaged with Warp [29]. The tilt-images containing all frames were sorted to generate the tilt-ordered raw tilt-series using a custom script and aligned with TomoJ v2.28 [30] using speeded up robust features (SURF) as fiducials.

## Acknowledgement

The authors acknowledge the use of instruments and assistance at the Monash Ramaciotti Centre for Cryo-Electron Microscopy, a Node of Microscopy Australia. This research used equipment funded by Australian Research Council grants: FEI Helios Cryo FIBSEM - ARC LIEF (LE150100132) and Titan Krios - ARC LIEF (LE120100090). This research was in part funded by ARC DP200103637. This work has been made possible in part by CZI grant DAF2021-225399, grant DOI https://doi.org/10.37921/334038myxhsa (to GR) and grant 2025-366327 (5022) GB-1633987 (to PC & GR) from the Chan Zuckerberg Initiative DAF, an advised fund of Silicon Valley Community Foundation (funder DOI: 10.13039/100014989).

## APPENDIX A Cracks in lamellae and FEM simulations

